# Spatio-temporal analysis of Vaccinia virus infection and host response dynamics using single-cell transcriptomics and proteomics

**DOI:** 10.1101/2024.01.13.575413

**Authors:** Alejandro Matía, Frank McCarthy, Hunter Woosley, Vincent Turon-Lagot, Sebastian W. Platzer, Jonathan Liu, María M. Lorenzo, Michael Borja, Kavya Shetty, Juliane Winkler, Joshua E. Elias, Rafael Blasco, Carolina Arias, Marco Y. Hein

## Abstract

Poxviruses are a large group of DNA viruses with exclusively cytoplasmic life cycles and complex gene expression programs. A number of systems-level studies have analyzed bulk transcriptome and proteome changes upon poxvirus infection, but the cell-to-cell heterogeneity of the transcriptomic response, and the subcellular resolution of proteomic changes have remained unexplored.

Here, we measured single-cell transcriptomes of Vaccinia virus-infected populations of HeLa cells and immortalized human fibroblasts, resolving the cell-to-cell heterogeneity of infection dynamics and host responses within those cell populations. We further integrated our transcriptomic data with changes in the levels and subcellular localization of the host and viral proteome throughout the course of Vaccinia virus infection.

Our findings from single-cell RNA sequencing indicate conserved transcriptome changes independent of the cellular context, including widespread host shutoff, heightened expression of cellular transcripts implicated in stress responses, the rapid accumulation of viral transcripts, and the robust activation of antiviral pathways in bystander cells. While most host factors were co-regulated at the RNA and protein level, we identified a subset of factors where transcript and protein levels were discordant in infected cells; predominantly factors involved in transcriptional and post-transcriptional mRNA regulation. In addition, we detected the relocalization of several host proteins such as TENT4A, NLRC5, and TRIM5, to different cellular compartments in infected cells. Collectively, our comprehensive data provide spatial and temporal resolution of the cellular and viral transcriptomes and proteomes and offer a robust foundation for in-depth exploration of virus-host interactions in poxvirus-infected cells.

## INTRODUCTION

Poxviruses are a large family of double-stranded DNA viruses that include several important pathogens, such as Variola virus (the causative agent of eradicated Smallpox), Monkeypox virus, and Vaccinia virus (VV), considered to be the prototype Poxvirus^1^. One of the key features of Poxvirus biology is the complex life cycle that occurs entirely in the cytoplasm, requiring the virus to encode its own transcription and replication machinery^2,3^. Despite the autonomy conferred by these features, many steps of the viral life cycle require virus-host interactions, such as entry, transport, translation, or egress. Furthermore, many viral genes have immunomodulatory functions, aimed at subverting the pathogen recognition systems in the cell, as well as counteracting antiviral signaling and effectors (i.e. IFN-stimulated genes -ISGs-) when activated^4–7^.

The VV genome is ∼200 kb in size and encodes about 200 ORFs which contributes to its versatility and efficacy as a biomedical therapeutic platform^8^. The transcription of VV is a tightly controlled sequential process, where the viral RNA polymerase carries out the viral transcription, and its temporal regulation is mediated by specific promoters and transcription factors^9–12^. The first genes to be expressed are termed early genes, and are transcribed from within the virion core immediately after the viral entry^13^. After DNA replication, the intermediate and late genes are expressed, which require the presence of early and intermediate proteins, respectively^14–16^. Simultaneously, the virus takes over the cell’s mRNA synthesis capacity by depleting mRNAs through mechanisms like mRNA decapping^17,18^.

Previous studies have focused on both the viral and cellular transcriptome dynamics using different sequencing methods, delineating the activities of viral promoters and viral gene regulation^19–23^. More recently, single cell approaches have been used to study VV infection in human blood monocyte-derived dendritic cells and canine mammary carcinoma cells^24,25^. Single-cell transcriptomics (scRNAseq) allows the characterization of the viral infection process in individual cells, capturing both host and viral transcripts at high resolution. Studies with pooled cell populations are unable to detect cell-to-cell differences, which therefore are often ignored. In this study, we aimed to temporally and spatially resolve the regulation of virus and host factors and study the intracellular changes in protein abundance and localization upon infection. To this end, we analyzed the transcriptomes of both VV and the infected cell across multiple timepoints using scRNAseq. Our aim was to delineate the infection process with a single-cell-temporal perspective. Using this approach, we profiled the transcriptional kinetics of VV in high resolution, depicting the main viral infection trajectory and its relationship with the cellular response to infection.

To achieve a more comprehensive understanding of gene expression dynamics during VV infection, we further investigated global proteome changes and the translocation of proteins within infected cells. A prominent change in the proteome during VV infection is the downregulation of host proteins, known as host shutoff. This well described process is caused by VV as well as other viruses. In VV-infected cells, the reduction of host protein expression is achieved primarily by the depletion of cellular mRNAs by virus-mediated decapping^17^. In addition to this global mechanism, a more specific downregulation of cellular proteins occurs through interactions between host and viral proteomes. Previous studies have identified many host proteins downregulated during the infection, including TRIM5, IFIT proteins, and others^26^. To get a comprehensive view of these changes, we have integrated the viral and cellular transcriptome and proteome datasets, allowing us to detect regulation patterns that reveal new potential interactions between virus and host. Finally, our mapping of the subcellular relocation dynamics of the host proteome revealed the translocation of host proteins that may be involved in cellular responses during VV-infection. Overall, our study provides a high resolution resource for host and poxviral multi-omics analyses, uncovering potential proviral and antiviral factors.

## RESULTS

### Single-cell resolution of the transcriptional landscape in VV-infected human cells

To resolve the heterogeneity of cell states over the course of infection, we recorded a time-course of infection of the VV Western Reserve strain in HeLa cells over the first 24 hours post-infection (h.p.i.), roughly equivalent to 2 replication cycles. To capture a large variety of cell states of infection and bystander cell activation, we used asynchronous, staggered infection conditions with two multiplicities of infection (Low MOI = 0.2 and High MOI = 2.0, see Methods). To minimize the batch effect, we used MULTI-seq^27^ to multiplex the experimental conditions before pooling all cells for droplet-based single-cell RNAseq profiling (Fig. 1A). We then used UMAP to analyze the transcriptional profiles of infected and control cells (Fig. 1B, Supplementary Data 1)^28^.

**Figure 1.**
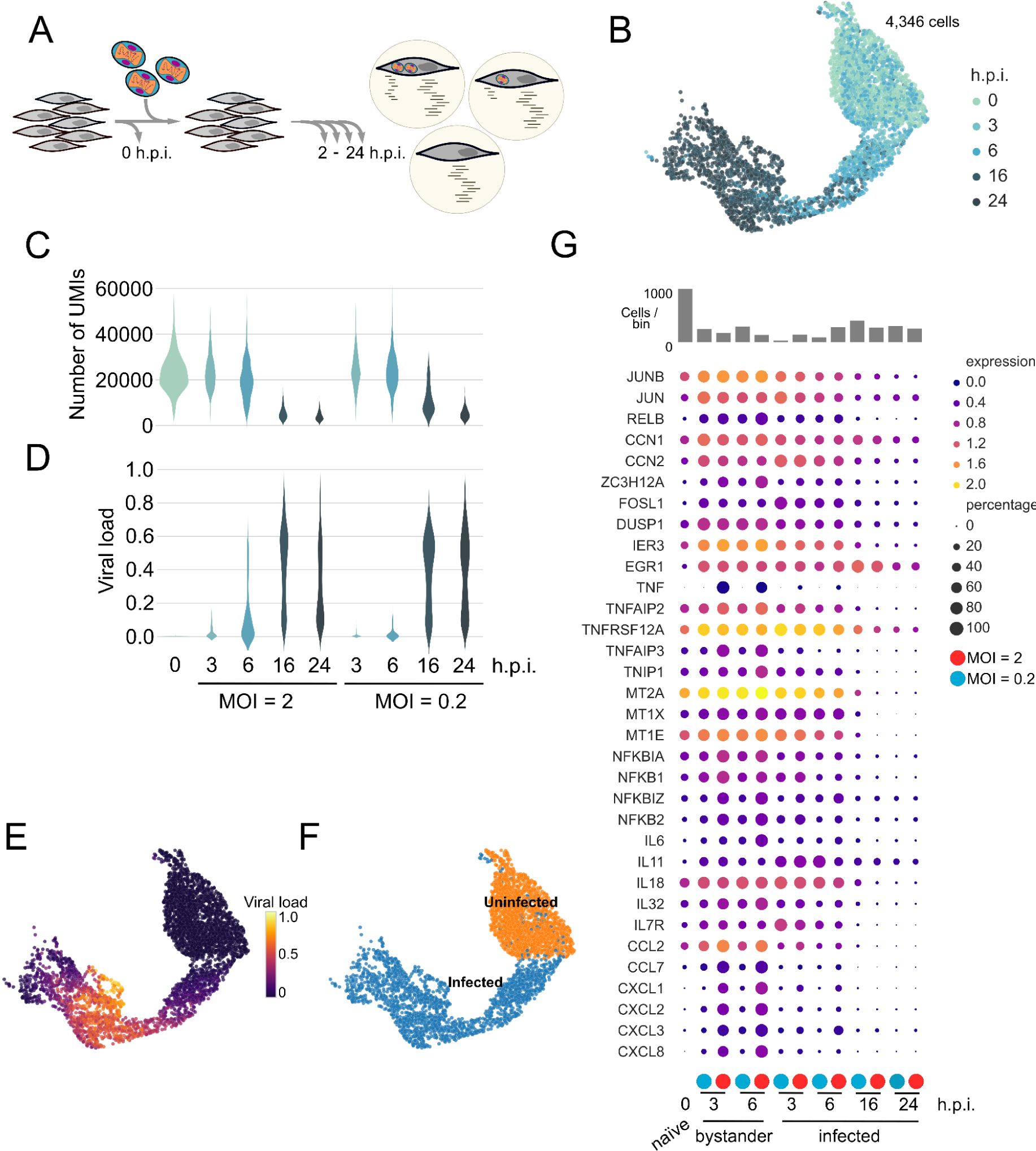
Single cell viral dynamics and host response (HeLa) A) Schematic representation of the experiment, where different MOIs and collection times were used. B) UMAP projection with cells colored by h.p.i. C-D) Host (C) and viral (D) dynamics evolution during the different experimental conditions. E-F) UMAP projections showing the viral infection trajectory (E) and the infection state classification (F). G) A selection from the top 100 differentially expressed genes (See Methods) between bystander vs naïve and infected vs naïve were selected to analyze the host response dynamics at different h.p.i.

Over the time of infection, we observed a reduction in the total number of unique molecular identifiers (UMIs) starting at 6 h.p.i., and dropping dramatically by 16 h.p.i. in cells infected at low and high MOIs, indicating the expected viral infection induced shutoff of cellular transcription (Fig. 1C). This phenomenon is well described and is the result of multi-level viral mechanisms leading to viral-induced host shutoff, where the host RNA and protein levels are modulated upstream and downstream of transcription^29^.

Concurrently with the onset of the transcriptional host shutoff, we observed a substantial increase in the viral transcriptome fraction (‘viral load’), consistent with previous bulk transcriptomics observations (Fig. 1D)^21^. Notably, the viral load followed a bimodal distribution in each condition, indicating that cells quickly transitioned from uninfected to infected state. In the early stages of infection, the viral load surged, reaching 40-60 % at 6 h.p.i. in some infected cells. This surge in viral load is a consequence of the significant reduction in host transcripts due to host shutoff, coupled with the rapid increase in viral transcription and accumulation of viral RNAs. As the infection progressed, the viral load peaked between 40 % and 90 % at 16 h.p.i., reflecting active viral transcription in the late phase. Intriguingly, at 24 h.p.i., viral loads appeared to drop, which we interpret as a consequence of the massive onset of cell death at that time. This observation is supported by increasing levels of mitochondrial mRNA levels with the progression of the infection, a common indicator of dying cells (Fig. S1). At late stages of infection (∼40 % of viral load), host shutoff reaches a maximum, and further increases in viral loads are independent of the reduction in human RNA counts, and therefore, only attributable to the continuous rise in viral counts (Fig. S2).

In UMAP projections, cells were arranged in order of increasing mitochondrial fraction, and concomitant increase of viral load, defining a trajectory of infection in the transcriptome space (Fig. 1E and Fig. S1). The trajectory culminates in cells with high mitochondrial fractions and viral loads that have regressed slightly past the peak (Fig. S1). We confirmed the temporal trajectory of infection by unbiasedly computing the trajectory inference using functional clusters obtained with Leiden algorithm^30,31^ (Fig. S3A and S3C).

We observed a wide distribution of viral load across individual cells at each time point of infection indicating that uninfected and several degrees of infected cells are present in each sample (Fig. 1C). To better differentiate the infected and uninfected cells we developed a classifier of infected cell state, based primarily on the number of viral counts (see Methods; Fig. 1F). We performed a gene expression analysis between relevant subpopulations of cells; naïve cells -control cells from the 0 h.p.i. condition-, infected cells, and bystander cells, defined as uninfected cells present in the infected samples (Fig. 1G).

Gene expression analyses showed that cellular stress response pathway genes were upregulated in both infected and bystander subpopulations. Many of these genes were expressed by the bystander cells at higher levels as compared to infected cells, suggesting possible paracrine signaling in response to infection (i.e. *JUN*, *JUNB*, *RELB*, *CCN1*, *DUSP1* and *IER3*). In bystander cells, we detected the consistent upregulation of *DUSP1*, which encodes a stress response protein acting as a restriction factor for several viruses, including VV^32^. Similarly, we detected the upregulation of genes involved in TNF and NF-κB inflammatory pathways, as well as many proinflammatory interleukins and chemokines consistent with previous reports^24^. Interestingly, we found distinct expression patterns for these transcripts, with some elevated in infected cells (i.e. *IL11* and *IL7R*), while others were detected at higher levels in the bystander population (i.e. *IL32* and *CCL2*). However, we did not detect the overexpression of interferon (IFN) and IFN-stimulated genes (ISGs) in these populations, likely a consequence of virus-mediated repression at the protein level^6^. Previous studies have shown that cytokine production in virus-infected populations is often confined to small subpopulations of abortively infected cells^33–35^. Clustering cells purely based on their transcriptome patterns indeed revealed that stress-related and cytokine transcripts are concentrated in a small cluster of infected cells (Fig. S3A, S3B, cluster i).

### Vaccinia virus infection triggers a default transcriptional program independent of the cellular background

To explore the conservation of the transcriptional responses we observed in HeLa cells in an alternative cell background, we used hTERT immortalized human fibroblasts (BJ-5ta) as an infection model. Based on the findings from HeLa cells, we added several experimental time points during the first 12 hours of infection, to better resolve the transition from early to late stages of infection (Fig. 2A, Supplementary Data 1). Similarly to the infection in HeLa cells, we observed the expected landmarks of VV infection in BJ-5ta, including the rapid transition of cells between the uninfected and infected states, and the rapid degradation of the cellular mRNAs starting at 6 h.p.i. (MOI = 2) (Fig. 2B).

**Figure 2.**
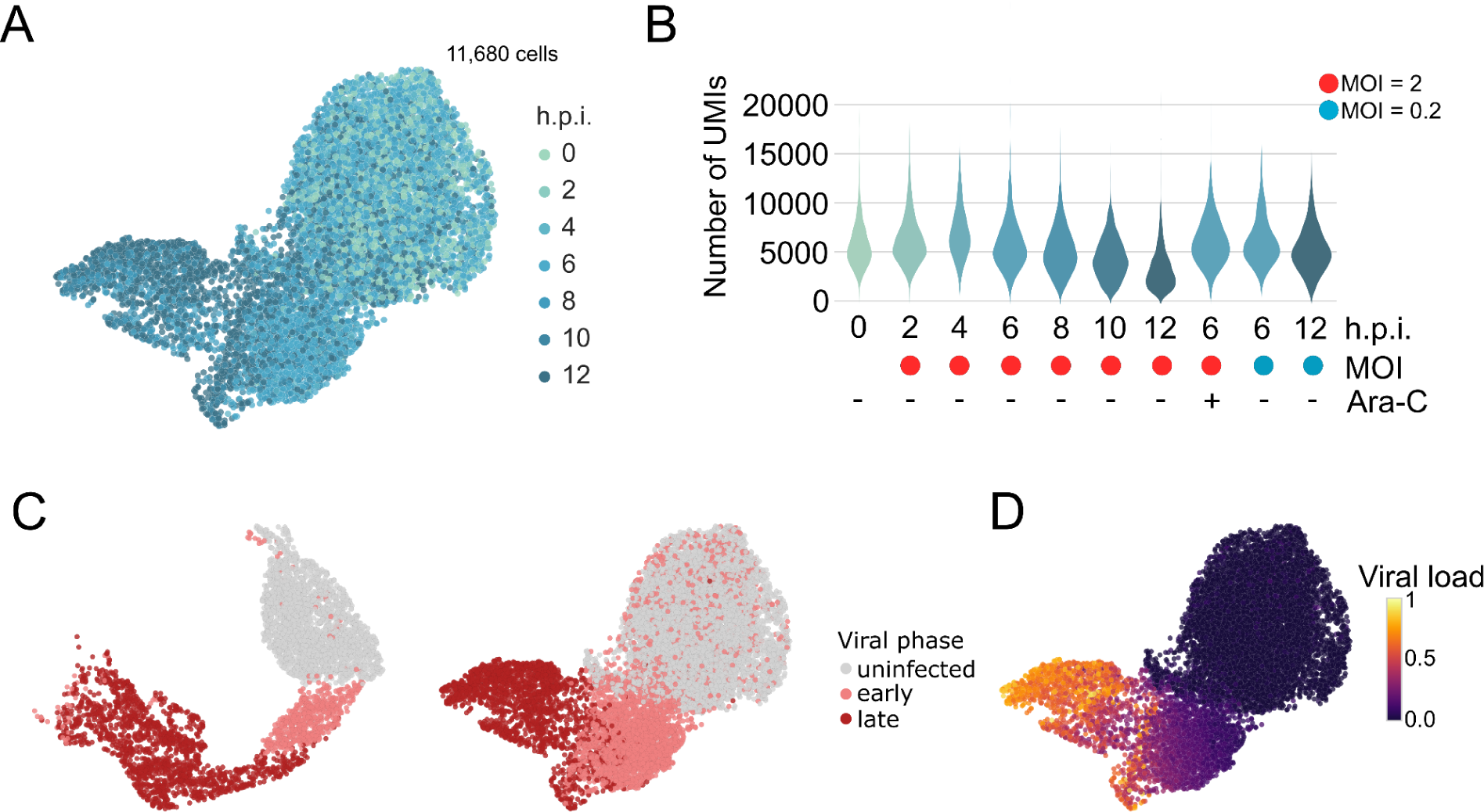
Host dynamics in human-derived fibroblasts (BJ-5ta) A) UMAP projection of BJ-5ta experiment with cells colored by h.p.i.. B) Host dynamics evolution during the different experimental conditions. C-D) UMAP projections showing the infection phase classification in HeLa -left- and BJ-5ta -right- (C) and the viral load (D).

A comparison of the host expression profiles in HeLa and BJ-5ta cells in the late phase of infection showed a generally conserved transcriptional response in those cell lines (Fig. S4). We only detected a few differentially expressed genes in the two cell lines at late infection. Genes like *TMEM107*, *CCN1*, *CCN2* or *TBX2*, displayed either a higher upregulation or a lesser degree of downregulation in HeLa cells compared to BJ-5ta. We found multiple histone-coding genes to be upregulated in BJ-5ta, which were downregulated in HeLa cells. Additionally, we also found many genes not expressed in one cell line that were strongly downregulated in the other. The differential depletion of most genes between the two cell lines may be explained by the differences in initial transcript abundances in the two cell lines.

Similar to our observations for the host transcriptome, a side-by-side comparison of the viral transcriptome kinetics in HeLa and BJ-5ta revealed the concordance in the timing of viral gene expression between these two cell types (Fig. 3). Overall, the viral gene expression profile showed the distribution seen in previous studies with many late (essential) genes enriched in the central portion of the genome^19–21^. The remarkable similarity of the viral gene expression patterns in HeLa and BJ-5ta, highlights the technical quality and reproducibility of our assay, and indicates that VV infection triggers a cell-type-independent viral expression program. Moreover, the resemblance in the overall infection trajectory in both cell types suggests that the limited number of genes with cell-line specific expression patterns does not affect the overall infection dynamics.

**Figure 3.**
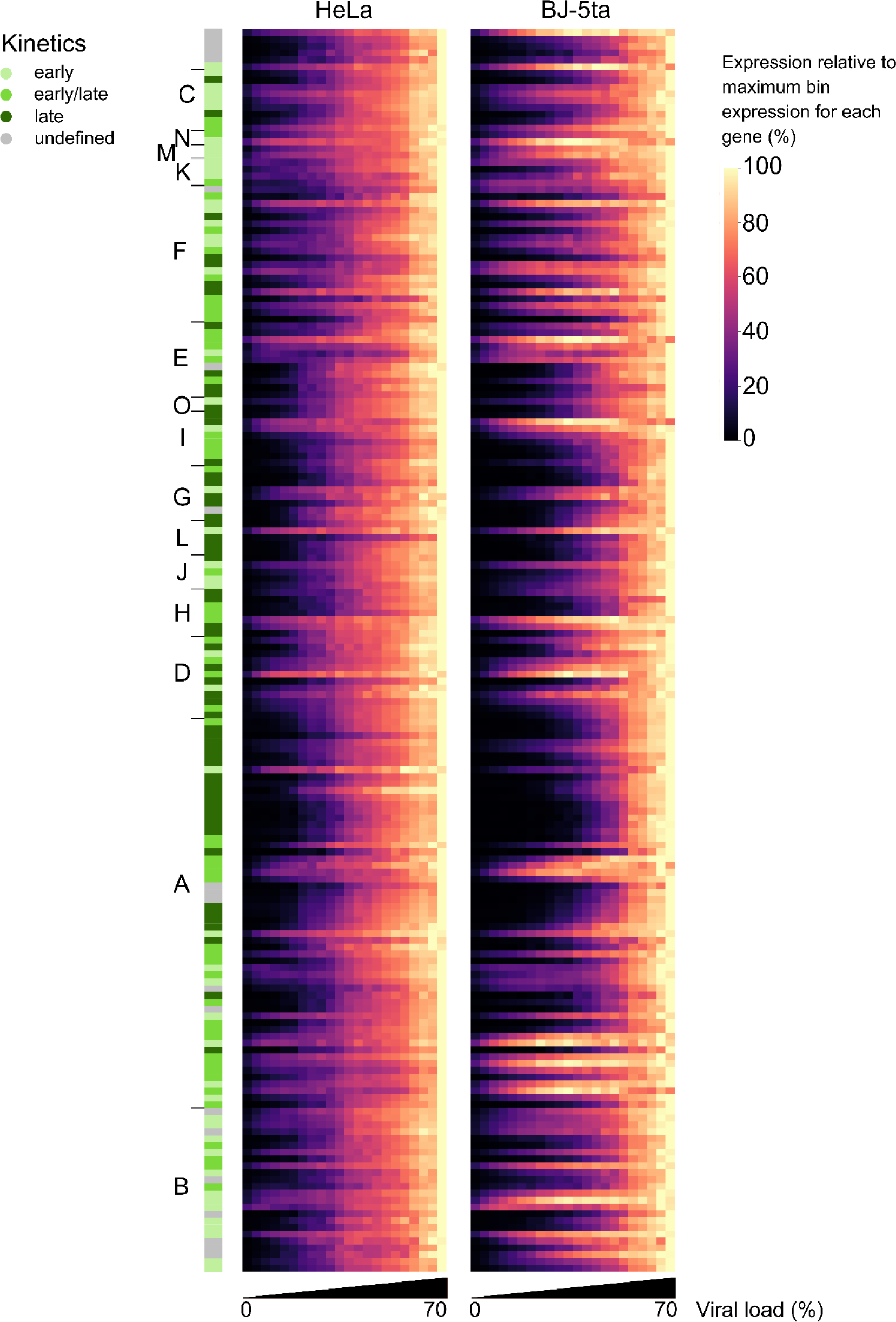
Viral dynamics at transcriptome level. Viral gene expression was represented as a function of the viral load (x-axis) for the ∼200 viral genes (ordered by genome position). Gene expression level in each bin was internally normalized to the maximum expression of that gene. Viral kinetic classes were also represented according to literature^19^.

Infected cells span a wide spectrum of transcriptional states. To distinguish between the pre-replicative (early) and post-replicative (late) phases of infection, we defined an additional classifier by using phase-specific marker genes (see Methods; Fig. 2C). To test the reliability of our classifier, we analyzed a population of cells pre-treated with Arabinose C (Ara-C), an inhibitor of viral DNA replication that blocks the transition to late phase, confirming that almost all infected cells in that condition are classified as “early” (Fig. S5A and S5B)^36^.

In summary, we detected viral early and late genes to be expressed in a tightly controlled temporal cascade during the infection cycle. In addition, we found a good correlation between the viral load and the sequential expression from early to late viral genes, consistent with host shutoff and the accumulation of viral RNA (Fig. 1E and 2D). Lastly, we used a cell cycle classifier to determine infection-driven transcriptional alterations of the cell cycle (see Methods). We observed a shift towards a transcriptional signature resembling the G1 phase, which was associated with late viral expression, similar to that reported during SARS-CoV-2 infection^35^ (Fig. S6).

### The host proteome is remodeled and relocalized during Vaccinia virus infection

Our transcriptomic analyses showed a pronounced remodeling of the host transcriptome in response to infection and highlights the organized kinetics of viral gene expression. To determine if similar changes are happening at the protein level, we captured the host and viral proteomes from HeLa cells at 16 h.p.i. at MOI = 2, corresponding to a robustly infected population prior to the onset of widespread cell death (see Fig. 1D). To this end, we prepared whole cell extracts, as well as crude subcellular fractions (corresponding to nuclear, organellar, and cytosolic proteins, respectively) from both infected cells and uninfected control samples, and analyzed the proteomes by label-free quantitative mass spectrometry.

In total, we quantified 8,382 proteins (8,203 human and 179 viral; Supplementary Data 2), several of which are modulated in VV infected cells. In accordance with previous reports, numerous host proteins were downregulated^37^ as a consequence of pre- and post-translational regulation, including IFN-related proteins such as the IFITs^38^ and IFNGR1, inhibitors of host translation such as EIF4EBP2 and EIF2AK1, and cell cycle checkpoint proteins such as p21 (*CDKN1A* gene) and p57 (*CDKN1C* gene) whose elimination leads to S and G2 phase accumulation (Fig. 4A)^39^.

**Fig. 4.**
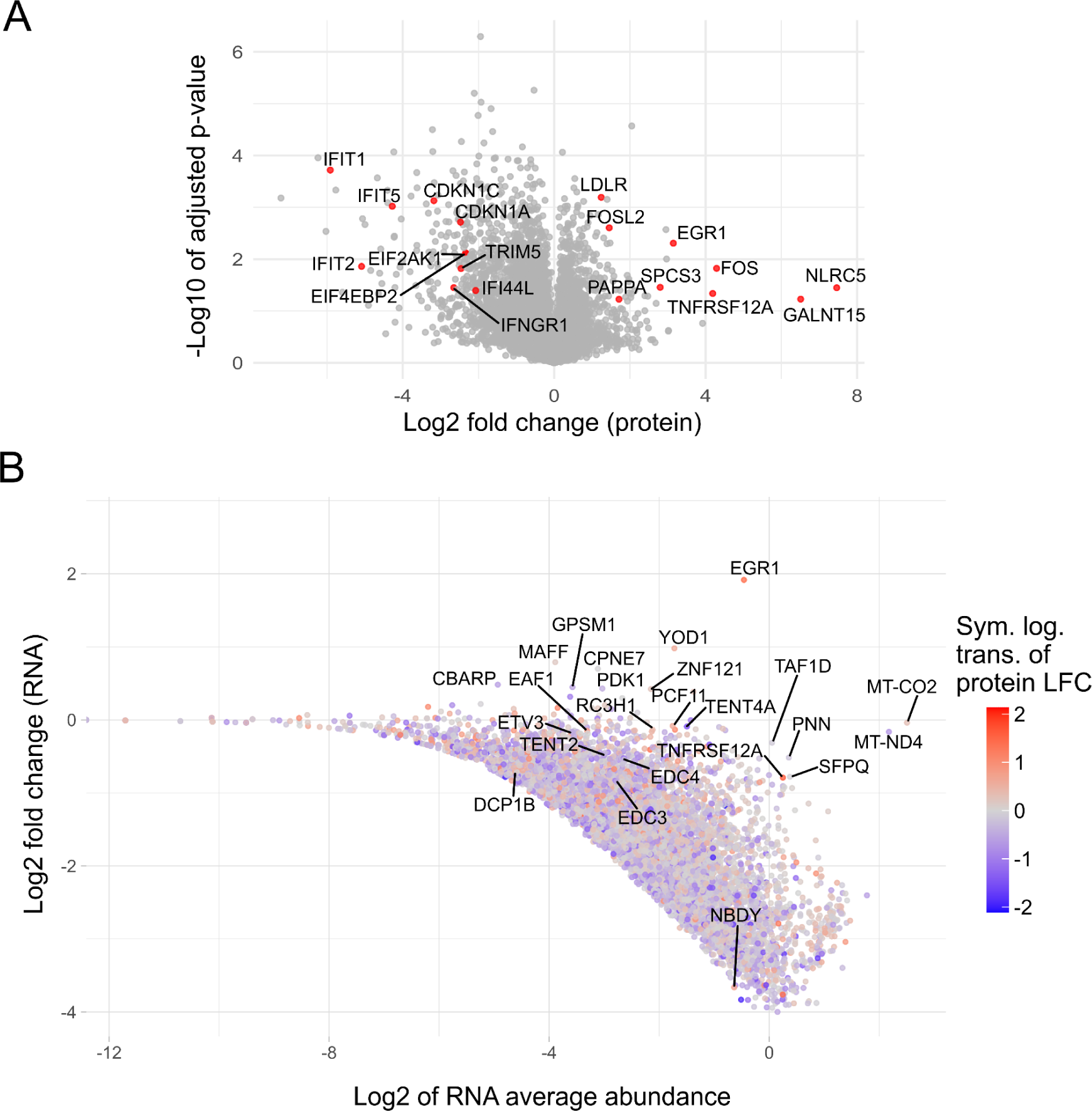
Host proteomic changes during the infection. A) Volcano plot representing the protein LFC of uninfected vs infected cells proteomes. Proteins discussed in the text were labeled. B) MA-plot representing the mRNA LFC as a function of the RNA abundance (log2-transformed). The color represents the protein LFC in a symmetric logarithmic (log1p) scale transformation. Proteins discussed in the text as well as the most extreme outliers were labeled.

Interestingly, even in the context of the strong, virus-induced host shutoff, we detected a small group of host proteins increasing in abundance during infection. Among these, the strongest upregulation we detected was for NLRC5, a pattern recognition receptor shown to reduce NF-κB and interferon type I activation upon stimulation with the dsRNA analog poly I:C or viral infection^40,41^. A second group of proteins highly upregulated in VV infected cells are associated with stress responses and MAPK pathway, including FOS, FOSL2 and EGR1, all of which are important for viral replication^42,43^. Addtionally, we found upregulation of TNFRSF12A, previously shown to be upregulated at the cell surface during infection^44^, and upregulation of PAPPA, which increases the availability of the Insulin-like growth factor, suggesting a strategic manipulation of cellular metabolism in favor of the virus^45^. Other proteins with significant upregulation were SPCS3 -a documented proviral factor for many viruses^46^-, GALNT15 -responsible for O-glycosylation-, and LDLR -a known viral receptor-^47^.

Next, we explored the correlation between RNA and protein levels in infected HeLa cells (Fig. 4B). While most of the host transcripts and proteins are downregulated as a consequence of the global host shutoff, we identified a group of proteins for which both mRNA and protein were upregulated. Among these factors, we found the upregulation of *ETV3* and *RC3H1*, involved in transcriptional and mRNA stability regulation downstream of STAT3. ETV3 is a transcriptional repressor activated downstream the anti-inflammatory STAT3 pathway, known to repress ISGs and stress response pathways, suggesting a direct, cis-repression mechanism of host transcription^48–50^. In a similar way, Roquin-1 (*RC3H1*), upregulated by the STAT1 and STAT3 pathways, promotes mRNA degradation by polyA-removal of transcripts harboring the constitutive decay element (CDE), such as *IER3*, *NFKBID*, *NFKBIZ*, *PPP1R10*, *TNF* and *TNFRSF4* which modulate NF-kB pathway^51–54^. Roquin-1 is also known to modulate miRNA stability leading to mRNA silencing by interacting with Dicer and Ago proteins^55,56^. Furthermore, our recent findings, and supported by other studies, showed that *RC3H1* has a proviral role in VV infection^57–59^. Taken together, our observations suggest that VV manipulates the host’s anti-inflammatory mechanisms precisely and add polyA tail-mediated mRNA destabilization to the list of known VV-mediated mRNA depletion strategies.

Several other genes exhibited upregulation at both mRNA and protein levels, including EGR1, which is expressed in the presence of the Vaccinia virus Growth Factor (VGF) and known to be important for VV replication^42,60^. Furthermore, proteins involved in transcription regulation such as EAF1 and PCF11, as well as ubiquitination-related proteins like YOD1, displayed a similar trend. The role of some of those factors in infection remains to be determined.

A second group of interest is comprised by the factors for which mRNAs are increased -likely due to transcriptional overexpression and/or shutoff escape-while their corresponding proteins are downregulated (both relative to the whole transcriptome or proteome set respectively), suggesting an active counteracting by the virus at protein level. Of particular interest among these, is the Terminal Nucleotidyltransferase 4A (*TENT4A* or *PAPD7*), which enhances cellular mRNA stability by non-canonical polyadenylation^61,62^. This process is facilitated by adaptor proteins such as ZCCHC14 and ZCCHC2 (the latter also found downregulated in our proteomics dataset) which recognize secondary structures in mRNA and bridge the interaction with these non-canonical poly(A) polymerases. Paradoxically, both TENT4A, its paralogue TENT4B, and their adaptor proteins stabilize viral mRNAs, and act as proviral factors for Hepatitis A and B viruses, Human Cytomegalovirus, and picornaviruses^63–66^.

Similar to TENT4, we observed the downregulation of the Terminal Nucleotidyltransferase 2 protein, also known as GLD-2 protein (*TENT2* or *PAPD4*), despite detecting elevated mRNA levels during infection. While TENT2 is also involved in mRNA regulation, it plays a relevant role enhancing miRNAs stability through 3’ modifications such as adenylation, guanylation, and uridylation^67–70^. During VV infection, the depletion of numerous host miRNAs has been reported, which could be at least in part explained by the TENT2 downregulation^71,72^. Interestingly the knockdown of TENT4 and TENT2 showed an increased VV infection phenotype in screen experiments, suggesting an antiviral function for these factors in the context of VV infection^57^. Other functional pathways enriched in this unique gene expression profile during VV infection (high transcript/low protein) include “Processing of Capped Intron-Containing Pre-mRNA”, “Chromatin Modifying Enzymes”, “Transcriptional Regulation by TP53”, and “cellular traffic”.

Finally, we identified a small group of factors characterized by low levels of RNA and high levels of proteins. Noteworthy within this group was the upregulation of NBDY during VV-infection, which is a component of the mRNA decapping complex, playing an important role in mRNA destabilization^73^. Other members of this decapping complex include EDC3, EDC4 and DCP1B, which displayed a similar gene expression profile. It remains to be determined if the higher protein levels for these factors are due to enhanced protein synthesis or protein stabilization during VV infection.

To evaluate the proteome localization in both naïve and infected cells we used subcellular fractionation, following a method that enables segregation of the nuclear, cytosolic, and organellar compartments. Firstly, to validate the accuracy of the different subcellular fractions, we used well-known protein markers of these compartments for calibration (See Methods) (Fig. S7A). Our analysis of host proteome localization revealed a pattern characterized by the nuclear-organelle axis and the organelle-cytosolic axis (Fig. 5A). We also detected a tendency towards higher degradation of organellar proteins upon infection, whereas cytosolic proteins were generally more resistant to viral-mediated shutoff (Fig. S7B). In examining the viral proteome, we detected a higher concentration of viral proteins in the organellar fraction, where viral factories are typically located, irrespective of the phase of infection (Fig. 5B and Fig. S7C).

**Fig. 5.**
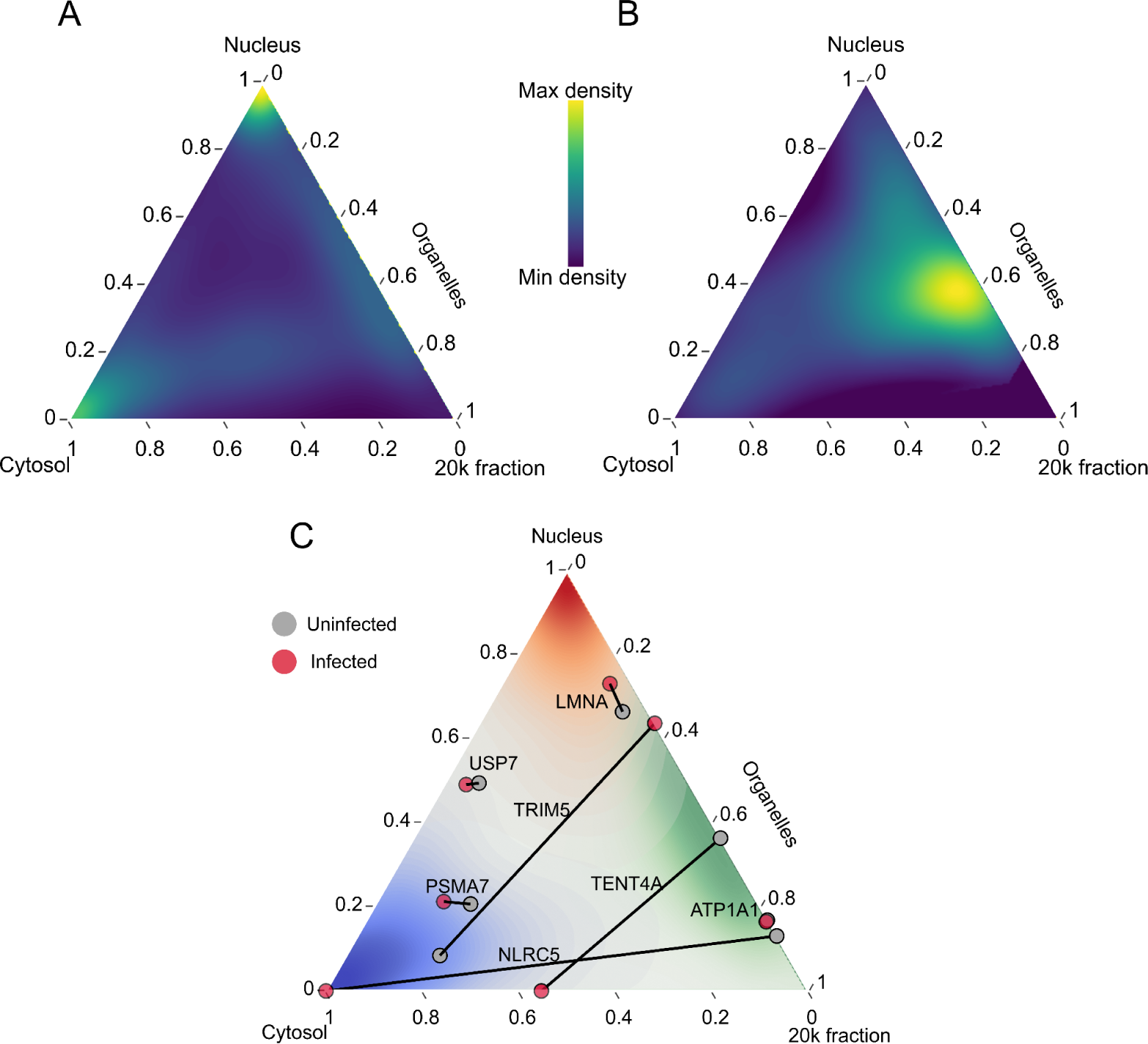
Viral and host proteome remodeling location during infection. A) and B) Density distribution of all host proteins (A) and all viral proteins (B) on a ternary graph of our subcellular fractions shows that viral proteins are predominantly located in organelle-like compartments, a distribution that differs strikingly from host proteins. C) Ternary plot representing the host proteome location during infection distributed in the 3 fractions (Nuclear markers in red, Cytosolic markers in blue, Organellar markers in green). A set of proteins was labeled for comparing the location before the infection (gray) and after (red). Control proteins with a stable location pattern during infection were also depicted: Lamin A/C (LMNA) for nucleus, Ubiquitin Specific Peptidase 7 (USP7) and Proteasome 20S Subunit Alpha 7 (PSMA7) for cytosol and nucleus (partially), ATPase Na+/K+ Transporting Subunit Alpha 1 (ATP1A1) for plasma membrane / organelles.

A comparison of the subcellular localization of host proteins between infected and uninfected cells revealed significant changes in the localization pattern of many proteins (Fig. 5C). Some examples are the antiviral factors TENT4A and NLRC5, which relocated from the organelle to the cytosolic compartment in infected cells, and TRIM5, typically found in the cytosol, which relocated to viral factories during infection^74^.

We also evaluated the location of known viral-host protein complexes and consistently observed the proximity of interacting partners, even when not all of the protein pool is in the complex (Fig. S7D). Our findings contribute to the understanding of the proteome dynamics in VV-infected cells and provide a valuable resource for spatial proteomics in the context of viral infections.

## DISCUSSION

The emergence of single-cell transcriptomics and advanced proteomics techniques has enhanced our understanding of the dynamics of viral infections and host responses. In this study, we have combined the power of these methods to gain a detailed and high-resolution picture of host and viral gene expression, alongside the localization and remodeling of the host and viral proteome during poxvirus infection, using Vaccinia virus as a prototypical model. VV rapidly alters host gene expression dynamics to take over the cell while the host fights back, activating innate immune responses to gain control over the infection.

One of the main events observed during the infection is a dramatic host-shutoff, which manifests primarily in a dramatic drop in the observed cellular RNA content. The rapid downregulation of cellular transcription is attributed both to pre-transcriptional (H3K9 and H4K20 methylation) and post-transcriptional events (RNA degradation)^29,75,76^. The substantial downregulation of host transcripts and host protein production becomes more pronounced as the viral cycle progresses. This effect is primarily executed by viral proteins that promote mRNA decapping, as well as altered splicing^17,77^.

In addition to inducing the global downregulation of host gene expression, poxviruses specifically counteract viral recognition systems to prevent immune activation and cell death^4,6^. Our transcriptional analyses revealed significant variations in the expression pattern of immune transcripts throughout the course of infection in infected and bystander cells. Among the strongest responses was the activation of stress response pathways, characterized by overexpression of genes such as NF-κB and TNF pathway genes, *JUN*, *FOS* or *FOXO1*. While the overexpression of these genes was robust in early infected cells, host shutoff impacted their expression as infection progressed. These findings highlight that despite the rapid downregulation of host gene expression, the infected cells can initiate stress/inflammation programs. In some cases, the activation of these pathways can benefit viral replication. One notable example of this is the upregulation of EGR1, a transcription factor that promotes VV replication in both cultured cells and mouse models^60^, though the precise mechanism remains to be elucidated. Conversely, VV markedly suppressed the expression of the interferon system in both tested cell lines, a response that has been shown to be detrimental for VV^78,79^.

Another noteworthy observation from our study is that despite the expression of protective immune-related mRNAs in bystander cells, the virus successfully infected them nonetheless at late timepoints, similarly to findings in experiments with HCMV^34^. These results align with experiments demonstrating the ability of VV to infect interferon-pretreated cells^80^. Adaptation of VV to human cell lines can explain this efficiency avoiding cell immunity – even in primary-like cells such as BJ-5ta – and why a higher restriction has been found in other animal cell lines, where host restriction factors interactions can be weaker^81^.

We observed a consistent trajectory of infection and similar host responses in the two cell types we used. All infected cells followed the same course of infection, with only minor differences in the expression pattern of a few stress-related genes. VV has a wide host range and it is able to infect various cell types in culture and animals *in vivo*^82,83^. The extent to which different and more restricted viral trajectories are apparent in other cells less permissive for infection – such as Chinese Hamster Ovary (CHO)^81^ – remains to be determined.

One of the advantages of using scRNAseq in infected cells is the possibility of identifying cells undergoing distinct stages of infection within the same cell pool. In our studies in HeLa and BJ-5ta cells, we identified the sequential transcription from early to post-replicative (intermediate and late) viral gene expression. Simultaneously, we observed the progressive increase of cellular shutoff, resulting in higher viral loads in the late stages of the infection, with over 60 % of total RNA being of viral origin. The viral phase classifier we developed proved to be a reliable tool for categorizing cells into early or late phases, as confirmed by the accurate classification of cells treated with the viral DNA inhibitor Ara-C.

After classifying infected HeLa and BJ-5ta into distinct viral stages, we demonstrated the striking congruence of the viral expression patterns. This underscores the independence of VV from the host system, triggering a ‘default’ viral program and highlights the independence from the cell that poxviruses, unlike other viruses, possess^84^. Despite the consistency of our datasets and previous publications, it is important to recognize that the transcriptional profile is a snapshot of the cell state and that additional regulation strategies, independent of the regulation of RNA expression, can modulate VV protein synthesis.

To improve our understanding of protein synthesis control during infection, we conducted an in-depth analysis of the host and viral proteomes during infection. Our findings are consistent with previous reports, and show a complex picture of viral-induced regulations at the protein level^26^. These data also reflect the effect of several VV proteins known to counteract antiviral host proteins by blocking their function or by marking them for degradation. Among the downregulated proteins, we observed the depletion of proteins associated with the interferon response system, such as IFIT proteins and IFNGR1. Previous studies showed that B8, a VV-expressed protein which structurally resembles IFNGR1, hijacks IFN-γ preventing the activation of its host receptor and the IFN signaling^85,86^. In addition, our analyses also show that IFNGR1 protein levels are reduced during infection, suggesting the existence of alternative viral mechanisms aimed at suppressing the activation of this interferon pathway.

We also noted a significant reduction in translation initiation inhibitors, including EIF4EBP2 and EIF2AK1. The downregulation of these factors, coupled with the sequestration of translation initiation factors within the replication compartments, highlight the strategies by which VV enhances a selective mRNA translation amid host shutoff^87,88^.

Other host proteins reduced during the infection include p21 (*CDKN1A*) and p57 (*CDKN1C*), both cell cycle checkpoint proteins whose attenuation promotes transition to G2 phase of the cell cycle^39^. This shift towards the G2 phase, and the degradation of p21 during VV infection have been well documented^89^. The arrest of the cell cycle in G2/M has been associated with the replication of other viruses including HIV and Sendai virus, and is linked to the downregulation of IFN-associated responses^90,91^. Surprisingly, transcriptome analyses showed a pattern of gene expression resembling that of G1 phase^89,92^. We hypothesize that this could be due to the RNA depletion masking the true transcript-level response or by the virus and host pulling in different directions – the cell at transcript level and the virus at a more effective, protein level.

To explore the connection between transcriptional regulation and protein synthesis in VV-infected cells, we integrated the scRNAseq and proteomics datasets. Our analyses show that for most host factors, the up or downregulation at both mRNA and protein levels align. Notably, we observed the relative upregulation (transcript and protein) for Roquin-1 (*RC3H1*), a pro-VV factor previously identified in our genetic screens^58^. RC3H1 is an E3 ubiquitin ligase with a RING domain and an RNA-binding CCCH zinc finger, involved in immune homeostasis and in promoting immune-relevant mRNAs downregulation through polyA removal^51–54,93,94^. Thus, we argue that RC3H1 upregulation might be enhancing the viral-mediated host shutoff. Our observations open the possibility for a new RC3H1-dependent viral strategy to control mRNA stability in VV-infected cells, which will be examined in future studies.

Interestingly, our integrative multi-omics analysis showed a notable discordance in the levels of mRNA and protein for many genes. This trend, previously documented in differentiating cells^95^, has not been reported during VV infection. This observation raises the possibility of delayed translation dynamics for specific mRNAs, and the stabilization or destabilization of specific host factors at the protein level. Poxviruses, including VV, encode E3 ubiquitin ligases and adaptor proteins known to target immune factors, including NF-kB, MHC-I, and CD4^6,96,97^. However, most of the targets for these proteins remain unknown. Future studies will help us understand whether some of the proteins downregulated in our proteome data are novel targets for these viral factors.

Among the factors displaying high mRNA and low protein levels, we found TENT4 and TENT2. We speculate that these factors are targeted for degradation by VV at protein level, despite their overexpression and escape of shutoff. Host shutoff escape may occur from a lesser affinity of the viral decapping enzyme D10 in transcripts containing few introns, a mechanism used for the escape of viral transcripts^98^. However, TENT4A gene contains 13 exons, which argues in favor of an upregulation as a consequence of gene overexpression. Therefore this seems to be an example of the cell and the virus fighting in opposing directions.

During VV infection, global miRNA levels are also depleted. A known mechanism behind this phenomena is the action of VP55, a viral encoded polyA polymerase that targets host miRNAs for degradation while protecting viral mRNAs^99^. TENT2 is known to stabilize certain miRNAs, and the downregulation of this protein in our dataset suggests another viral mechanism for the miRNAs overall levels depletion during infection.

Additionally, we found the Negative Regulator Of P-Body Association protein (*NBDY*) to be strongly downregulated at mRNA level, in opposition to its increased protein levels during the infection. This protein is part of the mRNA decapping complex and promotes mRNA destabilization, explaining why its upregulation during the infection might be beneficial for VV^73^. Here we showed the robustness of our approach for detecting potential proviral or antiviral factors, as well as uncovering novel viral-host interactions.

Finally, along with the high resolution transcriptome maps, we have mapped the subcellular host and viral proteomes during infection. With this approach, we have been able to detect the relocalization of several host proteins along with the infection, such as TENT4A, NLRC5 and TRIM5. These data suggest specific viral hijacking mechanisms for repurposing host protein location and function, providing novel angles at which the viral machinery can be counteracted.

## METHODS

### Cells

HeLa (ATCC CCL-2) and BJ-5ta (ATCC CRL-4001) cells were grown in Dulbecco’s Modified Eagle Medium (DMEM) + GlutaMAX-I (Gibco) supplemented with 0.1 mg/mL penicillin, 0.1 mg/mL streptomycin, and 10 % fetal bovine serum (FBS). BS-C-1 cells (ATCC CCL-26) were grown in Dulbecco’s Modified Eagle Medium (DMEM) + GlutaMAX-I (Gibco) supplemented with 0.1 mg/mL penicillin, 0.1 mg/mL streptomycin, and 5 % fetal bovine serum (FBS). All cells were grown in a 5 % CO2 incubator at 37 °C.

### Virus

Vaccinia virus WR strain (ATCC VR-1354) was used for all the experiments. Virus was grown in HeLa cells and supernatants from 48 h.p.i. were collected, cleared by centrifugation (10 min at 600xg) and filtered by 0.45 μm filters, frozen and thawed 3 times and titrated in BS-C-1 cells following the standard VV protocol for plaque assay^100^. All virus infections were performed in DMEM media containing 2 % FBS. For the experimental infections, the viral stock was thawed, sonicated and the virus was mixed with 2 % FBS media to achieve final MOIs of 0.2 and 2.

### scRNAseq sample and library preparation

Cells from each sample were trypsinized and tagged with a different LMO following established protocol^27^. After tagging, cells from all the conditions were pooled together and introduced in one lane of the Chip G in the Chromium (10x Genomics). RT-PCR was performed according to protocol and cDNA was PCR-amplified for 11 extra cycles. Library preparation for Illumina sequencing was performed according to protocol. Libraries were sequenced on the NovaSeq platform (Illumina).

### Bioinformatic Analyses

cellranger v5.0.1 software was used to read-mapping using VV WR (LT966077) and hg19 human reference genomes respectively. Scanpy v1.9.1 was used to run scRNAseq data analysis. Cell doublets or cells marked as unassigned (multiple or no MULTI-seq barcode, respectively) were filtered out before downstream analysis. Normalization was applied only for specific purposes to avoid alteration of gene expression ratios due to shutoff.

### Data curation

While in other experimental setups it is desirable to place a filter threshold to remove all the reads associated to cell debris, due to the virus-induced transcriptional host-shutoff, filtering out cells with a low UMI count would discard many infected cells. Following the same principle, we did not normalize the data for the absolute analyses to the total number of UMIs per cell, since this would alter the transcript-to-transcript ratio in an artificial manner in cells with truly different biological UMI counts. Therefore, cells with UMI count per cell > 200 were kept for the first experiment. For the second experiment, a cutoff of UMIs/cell > 800 was applied, since less debris was observed, probably because the lack of > 16 h.p.i. cells is translated into a better cell integrity. Additional filtering was applied after cell clustering using UMAP and Leiden algorithms to remove uninfected cells with low UMI count and high mitochondrial fraction clustering together.

### Classifiers

#### -Cell state classifier

Cells with a viral count > 125 were considered as infected considering background noise in uninfected cells. Cells with less or equal as 125 viral counts, > 5000 human counts and a mitochondrial fraction < 0.15, were considered as uninfected. Unclassified cells harboring high mitochondrial fraction were filtered out.

#### -Cell cycle classifier

Cell cycle scores were calculated as previously described, including a score for G1 phase^101^. RNA expression was normalized to each respective cell total. A modification of the algorithm used to calculate cell cycle phases was done, including the calculation of G1 phase with a set of marker genes specific to this phase, instead of assigning to G1 phase the cells that could not be assigned to either S or G2M phases. Marker genes for G1 phase: *ORC1, SP1, TAF1, MTF1, SIX5, SMARCC2, HCFC1, TFAP2A, KLF4* and *ESRRA.* For S phase: *MCM5, PCNA, MCM2, MCM4, UNG, MCM6, CDCA7, DTL, PRIM1, UHRF1, RPA2, WDR76, SLBP, CCNE2, POLD3, MSH2, CDC6, TIPIN, CASP8AP2, POLA1, CHAF1B* and *BRIP1*. For G2M phase: *UBE2C, BIRC5, TPX2, TOP2A, NDC80, NUF2, MKI67, CENPF, TACC3, CCNB2, CKAP2L, AURKB, BUB1, KIF11, GTSE1, KIF20B, HJURP, CDCA3, CDC20, TTK, CDC25C, KIF2C, DLGAP5, CDCA2, CDCA8, KIF23, HMMR, AURKA, PSRC1, ANLN, CENPE, NEK2, GAS2L3* and *CENPA*. These marker genes were selected based on previous publications^102,103^.

#### -Viral phase classifier

Viral phase scores were calculated as previously described^101^, including early, late scores according to a set of viral marker genes. An initial list of genes was selected based on current bibliography^104,105^ and was then shortened keeping the genes that matched their peak expression with the assigned bibliographic classification.

The assignment to each viral phase was performed as follows: *If early score is lower than late score, or if late score is higher than 0.4, then it is ‘late’. If early score is lower than 0.15, then it is ‘uninfected’. Otherwise, it is ‘early’*. Marker genes for early phase: *C6L, C5L, N2L, M1L, M2L, K1L, K3L, K7R, F1L, VACVWR_00430, F11L, E5R, I3L, G5.5R, L2R, A8R, A29L, A31R, A40R, A44L, A51R, B12R, B13R* and *B19R*. Marker genes for late phase: *C8L, C3L, F3L, F9L, F13L, E8R, E10R, E11L, O2L, I1L, I8R, G3L, G4L, G7L, G9R, L1R, L3L, L4R, L5R, J1R, H2R, H7R, D2L, D3R, D8L, D13L, A2L, A2.5L, A6L, A7L, A9L, A10L, A11R, A12L, A13L, A14L, A14.5L, A15L, A16L, A19L, A21L, A26L, A27L, A28L, A30L, A38L, A45R, VACVWR_01470* and *VACVWR_00680*. During the late phase, many VV genes are transcribed as polycistronic RNA, which results in the presence of early RNA sequences in the cell. To avoid this limitation, we assigned the cells to the late phase only considering the late score. The threshold for early to late transition was selected using late phase-specific marker genes expression.

### Gene Expression analysis

To stabilize the variance of low-abundance genes, a pseudo-count adjustment (0.1) was applied before calculating the LFCs. Top differentially expressed genes were selected based on their LFC value. The top 100 was used for downstream analyses.

### Proteomics - Infections and Fractionation

HeLa cells were seeded in 10 cm dishes. Once they achieved a 80 % confluency, cells were infected with VV at MOI = 2 for 16 h, in the same way as the infections for single cell transcriptomics experiments. At 16 h.p.i., infected cells and uninfected control cells were washed with PBS and harvested and prepared for both whole-cell proteomics as well as subcellular fractionation exactly as described^106^. In brief, whole cell extracts were generated by scraping cells directly in SDS lysis buffer (5 % SDS, 50 mM triethylammonium bicarbonate (TEAB) pH 7.55). Subcellular fractions were generated by first homogenizing cells mechanically by resuspending cells in non-denaturing hypotonic buffer (25 mM Tris-HCl pH 7.5, 50 mM sucrose, 0.2 mM EGTA, 0.5 mM MgCl_2_, complemented with protease and phosphatase inhibitor cocktail (Thermo Fisher Scientific, #PI78443) and forcing them four times through a 23G blunt needle (SAI Infusion Technology #B23-100). After restoring osmolarity by adding 1/10 volume of 2.5 M sucrose, 0.2 mM EGTA, 0.5 mM MgCl_2_, subcellular fractions were generated by first centrifuging at 1,000 ×g for 10 min (4°C) to generate a nuclear pellet. The supernatant was further further centrifuged at 20,000 ×g for 45 min (4°C), resulting in a pellet containing mainly organellar proteins, and a soluble cytosolic supernatant. Both pellets were resuspended in SDS lysis buffer. The cytosolic supernatant was supplemented with 20 % SDS to yield 5 % SDS final. All samples were then boiled and sonicated. Downstream sample preparation and proteomics data acquisition was performed exactly as described^106^.

### Proteomics - Data Processing

For proteomics data processing, we used MaxQuant^107^ v. 2.2.0.0 with the match between runs feature enabled and MaxLFQ^108^ for label-free quantification (using 1 min. LFQ ratio count). Spectra were searched against human protein sequences (downloaded from Uniprot on 2021-07-17, canonical and isoform sequences) and a VV proteome reference (also downloaded from Uniprot; all entries with taxonomy ID 10254). A 1% false discovery rate was set at the peptide spectrum and protein matching with a minimum peptide length of 7, 2 missed cleavages were allowed, and enzyme was set to trypsin. Oxidation of methionine and protein N-terminal acetylation were set as variable modifications along with carbamidomethylation of cysteines set as a fixed modification.

MaxQuant output tables were further processed in Perseus^109^, using the Proteomic Ruler plugin for the estimation of protein copy numbers^108^ on the whole cell extract samples. We transformed protein LFQ intensities in the subcellular fractions as fractions of total intensity for visualization in ternary plots. A set of location marker proteins was used to establish the distribution pattern of Nuclear, Organellar or Cytosolic proteins in uninfected and infected conditions^110^. Ternary plots were built using the Plotly python library.

### Pathway analysis for transcriptomics + proteomics

We used a Generalized Additive Model (GAM) to establish a cutoff that reflects the profile of the transcriptomics distribution, based on RNA abundance and transcriptomics LFCs. Outliers above the curve were selected for downstream analysis. Reactome Pathway Analysis R package was used for obtaining the enriched pathways^111^.

## Supporting information

Supplementary Data 1: Single-cell transcriptomics metadata for VV-infected HeLa and BJ-5ta datasets.

Supplementary Data 2: Protein quantification (log2 LFQ intensities) in whole cell extracts and subcellular fractions of VV-infected HeLa cells.

## ACKNOWLEDGEMENTS

We want to thank Amy Kistler for helpful discussions and commenting on the manuscript. We thank Angela Detweiler, Sheryl Paul and Honey Mekonen for deep sequencing assistance. A.M. was the recipient of a predoctoral contract from Subprograma Estatal de Formación, Spain. We thank the Chan Zuckerberg Biohub and its donors Priscilla Chan and Mark Zuckerberg for funding this work.

## AUTHOR CONTRIBUTIONS

A.M., C.A., and M.Y.H. conceptualized the study. A.M., and M.Y.H. performed the single-cell and proteomics experiments. M.B. assisted with single-cell sample preparation. F.M. and J.E.E. performed mass spectrometry. H.W., V.T.L., S.W.P., and K.S. performed the validation experiments. A.M, and M.Y.H., analyzed the data with input from C.A. R.B., C.A., and M.Y.H. supervised the study. A.M., C.A., and M.Y.H. wrote the manuscript with input from J.W. All authors commented on the manuscript.

## SUPPLEMENTARY DATA

Supplementary Data 1:

Single-cell transcriptomics metadata for VV-infected HeLa and BJ-5ta datasets.

Supplementary Data 2:

Protein quantification (log2 LFQ intensities) in whole cell extracts and subcellular fractions of VV-infected HeLa cells.

## SUPPLEMENTARY FIGURES

**Figure S1.**
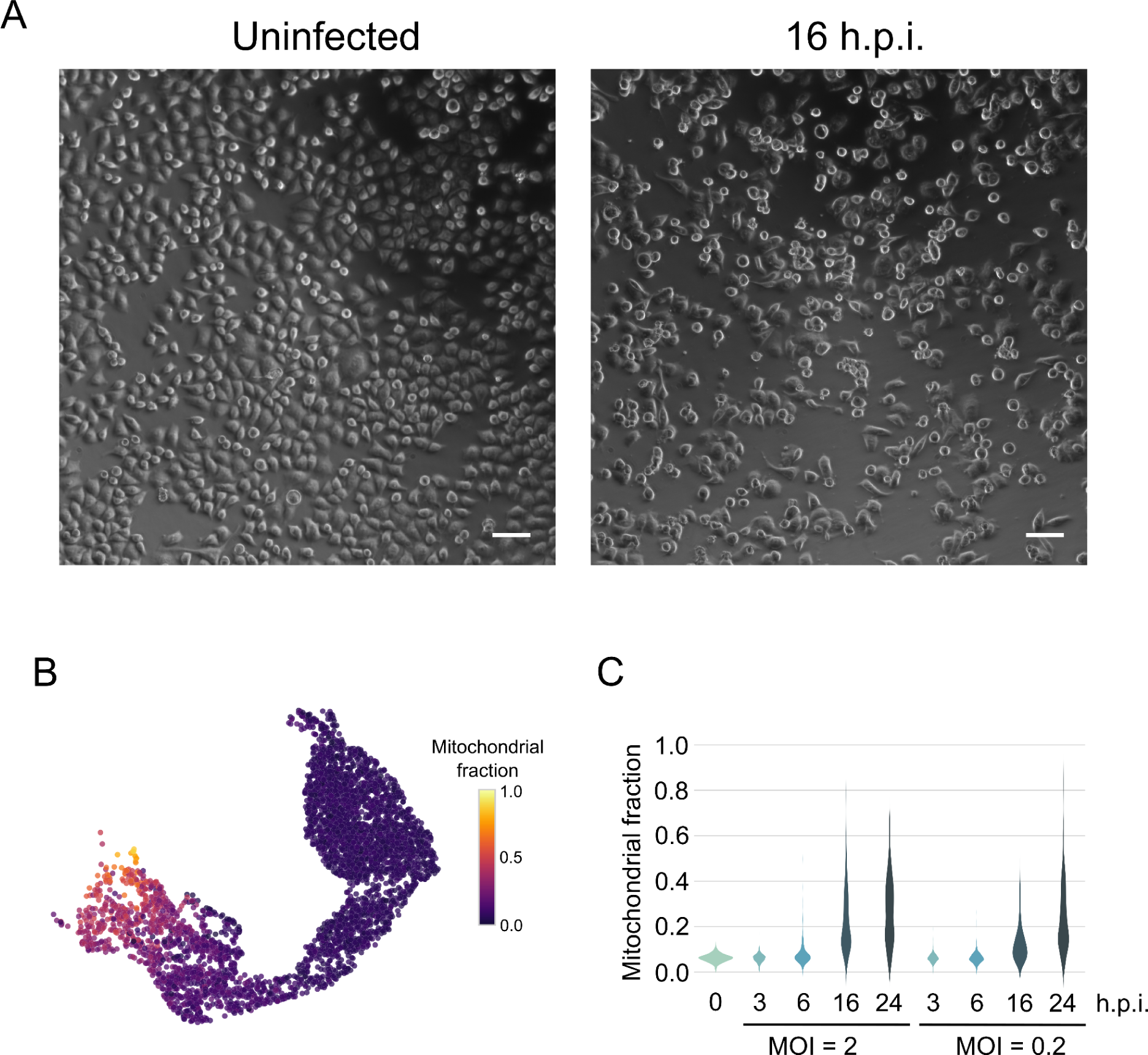
Viral infection cytopathic effect in human cells. A) Microscopy of HeLa cells (10X) uninfected or infected at MOI = 2 for 16 h with VV. Scale bar: 100 μm. B) UMAP projection of HeLa cells colored by mitochondrial fraction. C) Mitochondrial fraction was calculated for each cell and was represented as a function of the experimental condition.

**Figure S2.**
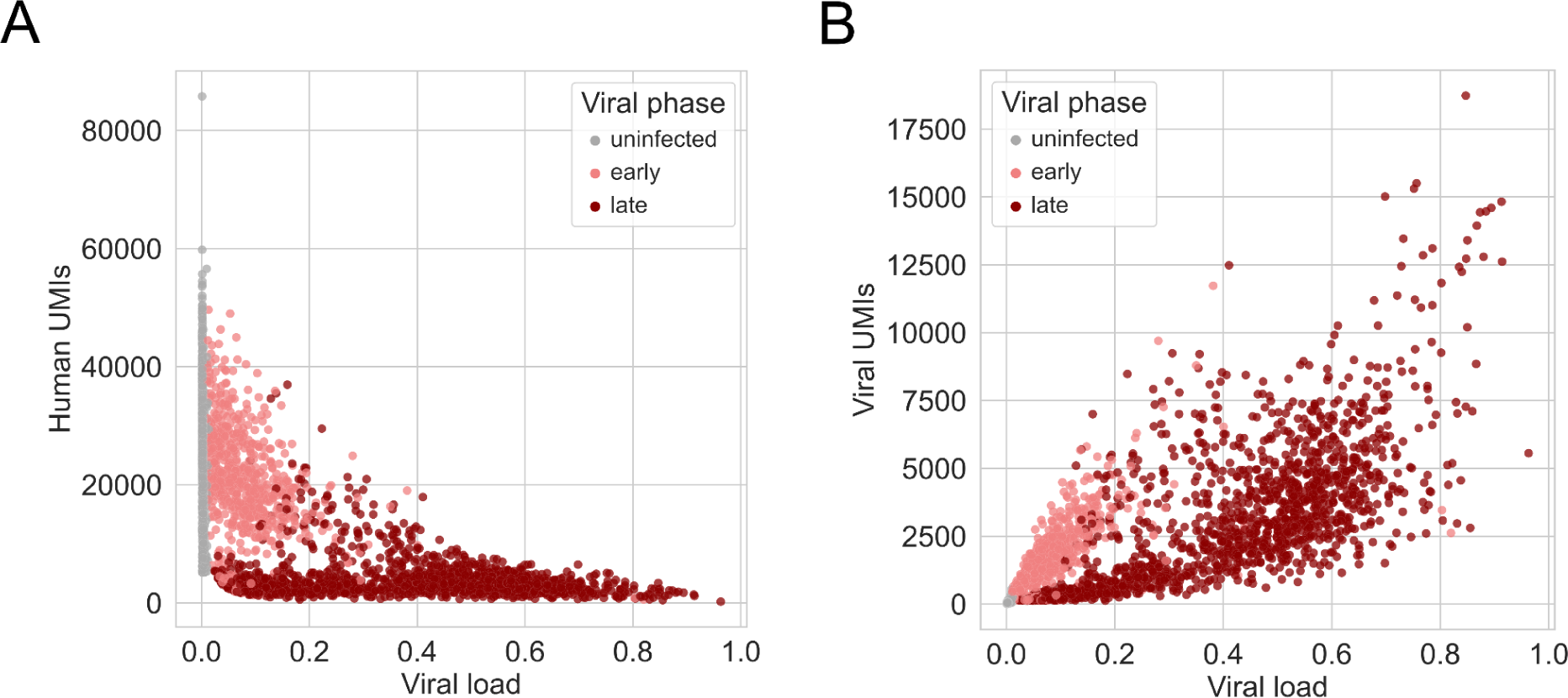
Host (HeLa cells) and viral dynamics as a function of the viral load. A) Host mRNA molecules (abundance quantified by UMIs) reduction is shown as the viral load (viral UMIs / total UMIs) increased. B) Viral UMIs increase in accordance with viral load differently in early or late-assigned cells due to concurrent shutoff in the latter.

**Figure S3.**
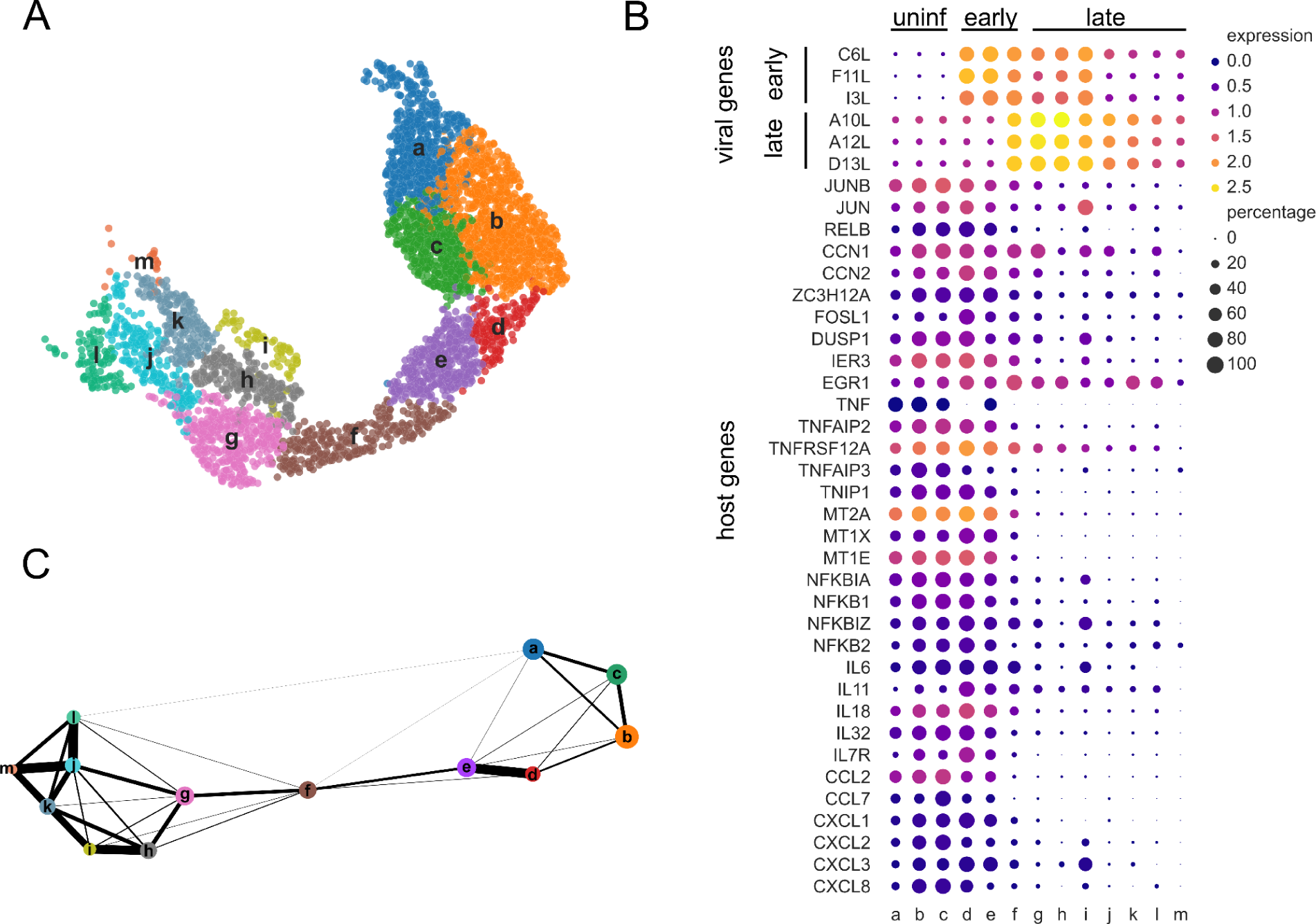
Clustering analysis of HeLa cells. A) UMAP projection colored by cluster-based classification. B) A set of the top differentially expressed genes between bystander vs naïve and infected vs naïve was selected to analyze the host response dynamics at different h.p.i.. Viral genes are also represented to follow the infection trajectory through the clusters. C) Partition-based graph abstraction (PAGA) representation of the inferred trajectories based on functional clusters^31^.

**Figure S4.**
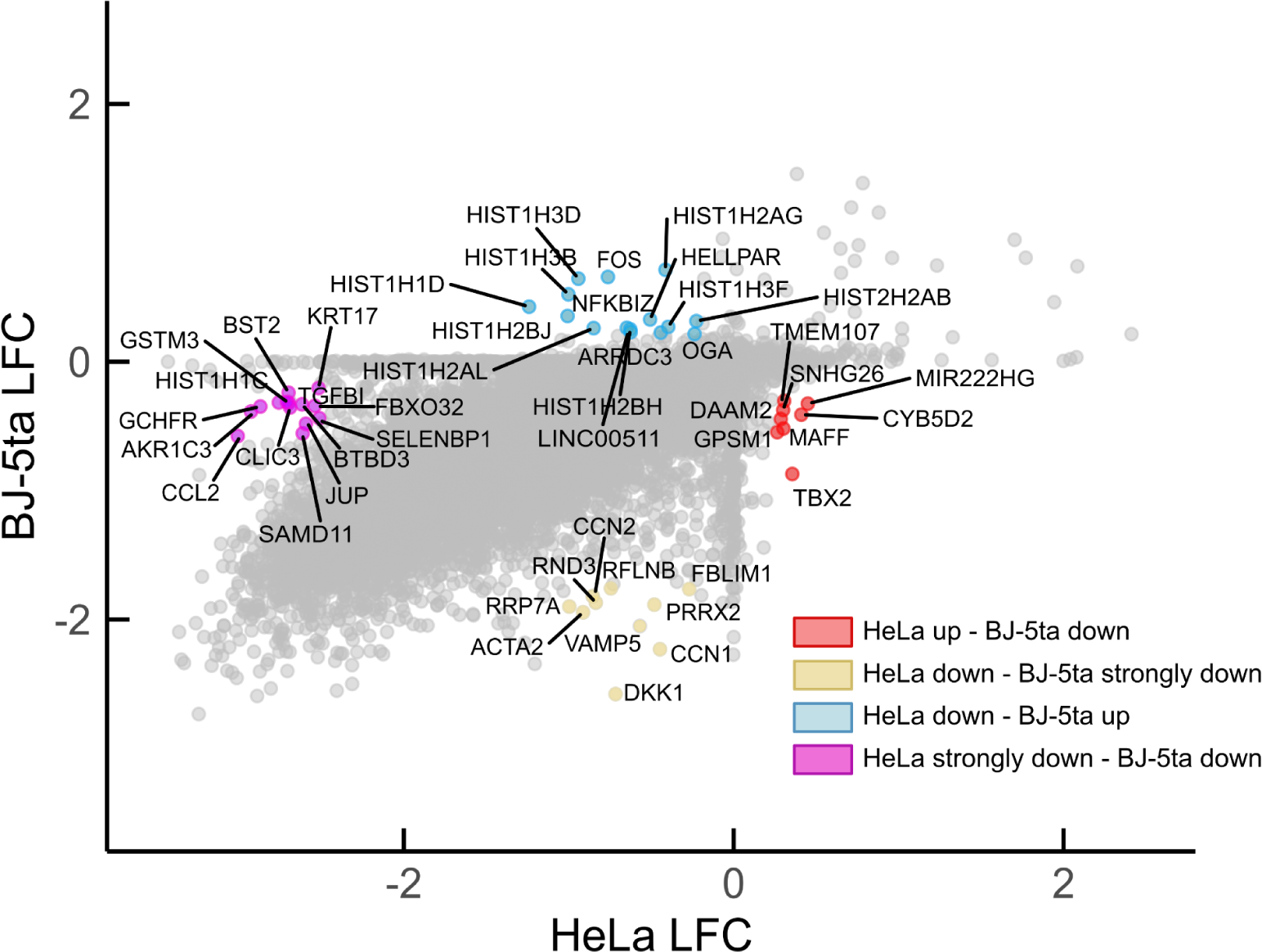
Side by side HeLa and BJ-5ta host response comparison. Log2 fold changes for each cell line were represented. The outliers from differentially relevant patterns were labeled, each color representing a different pattern.

**Figure S5.**
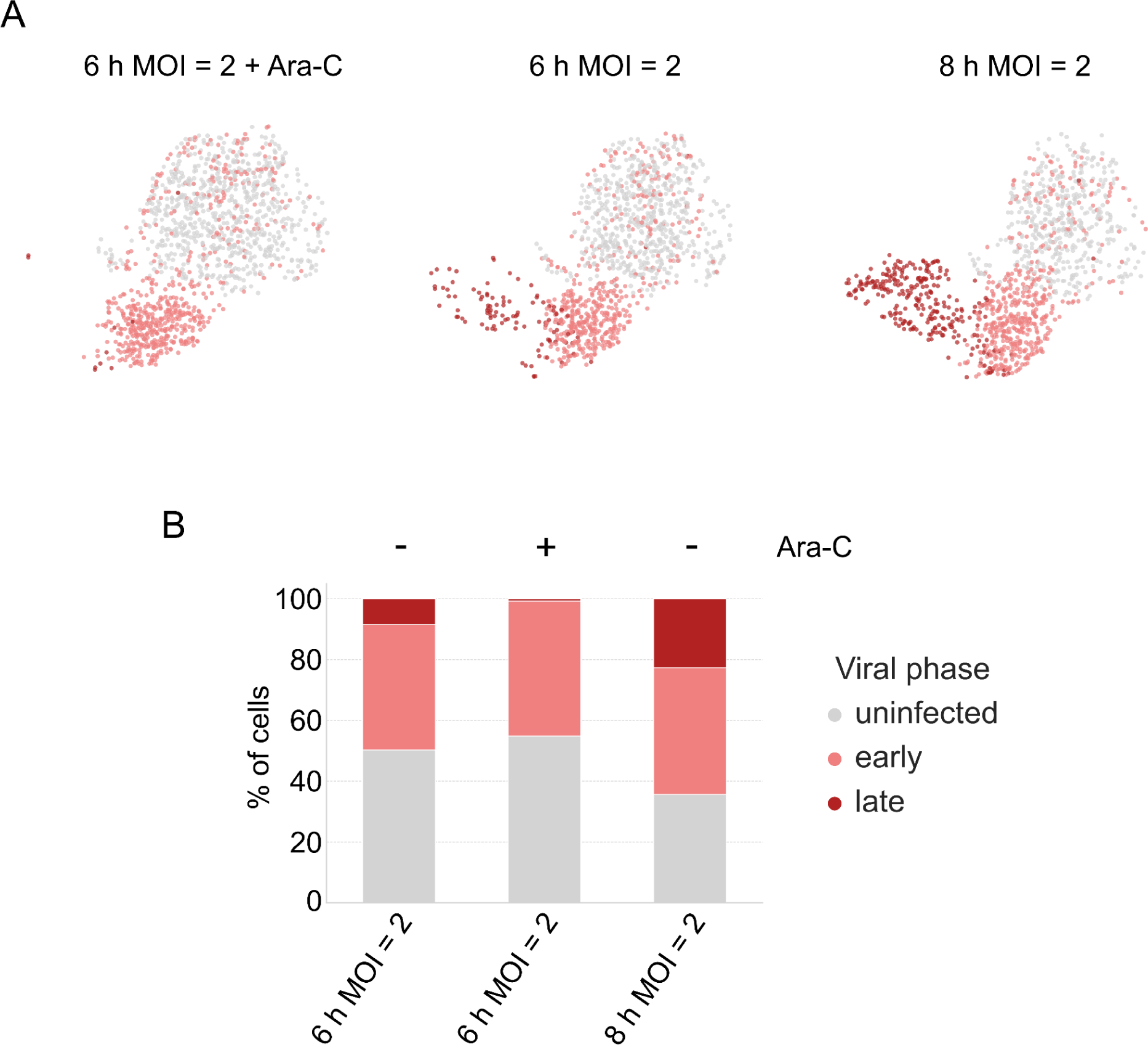
Effect of Ara-C at single cell level in human-derived fibroblasts. A) UMAP projections of BJ-5ta cells for each experimental sub-condition shows the effect of Ara-C inhibiting transition to late phase. B) The number of cells in each phase was quantified in each condition.

**Figure S6.**
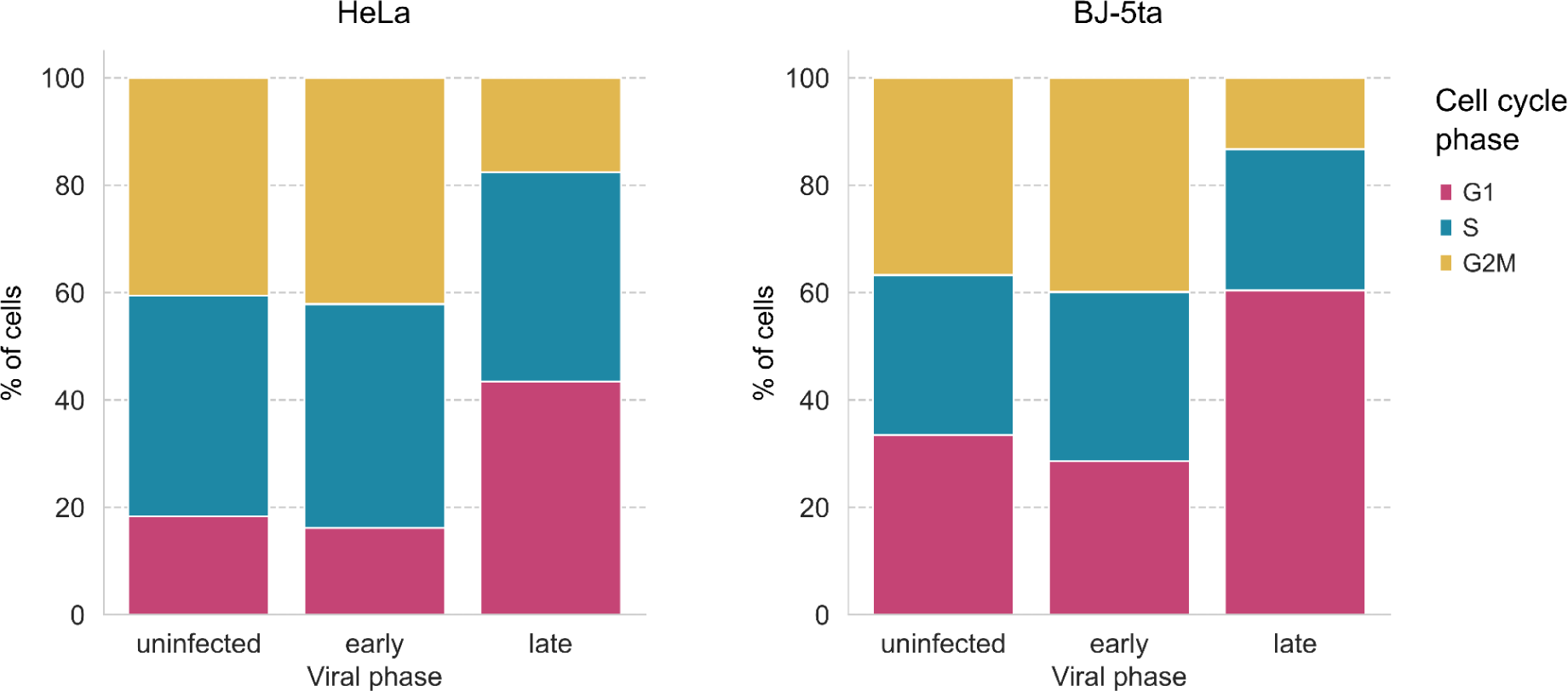
Cell cycle transcriptional signature during infection. Cells were assigned to G1, S or G2-M -like phases according to their transcriptional profile. The number of cells in each cell-cycle phase was quantified and represented for each phase of the viral cycle.

**Fig. S7.**
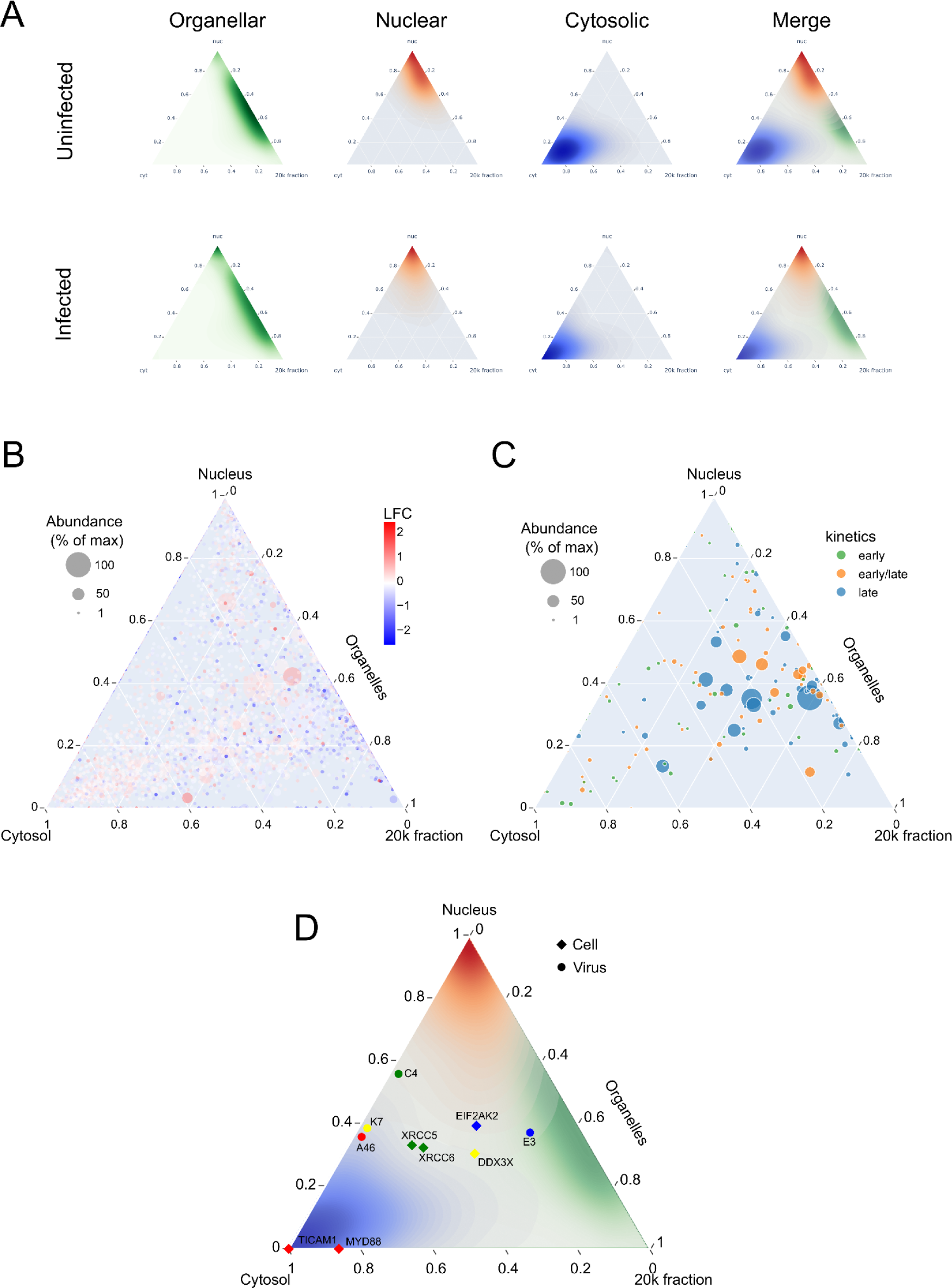
Proteomic distribution of host and virus during infection. A) Densities corresponding to each fraction were calculated and represented on the ternary plot for both uninfected and infected HeLa cells. B) Host proteome was represented, each dot being a protein, with the size corresponding to its relative abundance and the color corresponding to the protein LFC in the whole cell extract. C) Viral proteome was represented, each dot being a protein, with the size corresponding to its relative abundance and the color corresponding to its described gene expression kinetics. D) Host-virus interaction-complexes were overlaid on the ternary plot, each color corresponding to a known complex.

